# CRL5-dependent regulation of Arl4c and Arf6 controls hippocampal morphogenesis

**DOI:** 10.1101/2020.01.10.902221

**Authors:** Jisoo S. Han, Keiko Hino, Raenier V. Reyes, Cesar P. Canales, Adam M. Miltner, Yasmin Haddadi, Junqing Sun, Chao-Yin Chen, Anna La Torre, Sergi Simó

**Affiliations:** Department of Cell Biology and Human Anatomy, University of California Davis, CA 95616, USA; Department of Pharmacology, University of California Davis, CA 95616, USA; UC Davis Center for Neurosciences and Department of Psychiatry and Behavioral Sciences, University of California Davis, CA 95616, USA

## Abstract

The small GTPase Arl4c participates in the regulation of cell migration, cytoskeletal rearrangements, and vesicular trafficking in epithelial cells. The Arl4c signaling cascade starts by the recruitment of the Arf-GEF cytohesins to the plasma membrane, which in turn engage the small GTPase Arf6. In the nervous system, Arf6 regulates dendrite outgrowth in vitro and neuronal migration in the developing cortex. However, the role of Arl4c-cytohesin-Arf6 signaling during brain development and particularly during hippocampal development remain elusive. Here, we report that the E3 ubiquitin ligase Cullin 5/Rbx2 (CRL5) controls the stability of Arl4c and its signaling effectors to regulate hippocampal morphogenesis. Rbx2 knock out causes hippocampal pyramidal neuron mislocalization and formation of multiple apical dendrites. The same phenotypes were observed when Cullin 5 was knocked down in pyramidal neurons by in utero electroporation. We used quantitative mass spectrometry to show that Arl4c, Cytohesin-1/3, and Arf6 accumulate in the telencephalon when Rbx2 is absent. Arl4c expression is post-transcriptionally regulated, with a peak in expression at early postnatal stages, and is localized at the plasma membrane and on intracellular vesicles in hippocampal pyramidal neurons. Furthermore, we show that depletion of Arl4c rescues the phenotypes caused by Cullin 5 knock down in the hippocampus, whereas depletion of Arf6 exacerbates over-migration. Finally, we show that Arl4c and Arf6 are necessary for the dendritic outgrowth of pyramidal neurons to the most superficial strata of the hippocampus. Overall, we identified CRL5 as a key regulator of hippocampal development and uncovered Arl4c and Arf6 as novel CRL5-regulated signaling effectors that control pyramidal neuron migration and dendritogenesis.

## INTRODUCTION

Arl4c is a small GTPase that belongs to the <u>A</u>DP-<u>r</u>ibosylation factor (Arf)-like 4 protein sub-family (Arl4) (Matsumoto et al., 2017; Sztul et al., 2019). Despite the high level of Arl4c (also known as Arl7) mRNA detected in human brain tissue, its function in the nervous system remains unknown (Jacobs et al., 1999). In epithelial cells, Arl4 proteins participate in actin cytoskeleton rearrangement, vesicle dynamics, and cell migration (Matsumoto et al., 2017). In particular, Arl4c regulates filopodia formation and epithelial cell migration through Cdc42 activation (Chiang et al., 2017). In a different study, cultured intestinal epithelial cells treated with Wnt3a and EGF triggered the upregulation of Arl4c, promoting proliferation, cell migration, and tubulation by activating Rac1 and inhibiting RhoA (Matsumoto et al., 2014). Molecularly, Arl4 proteins, including Arl4c, can recruit the Arf-GEF cytohesins (Cyth-1/4) to the plasma membrane, which in turn can bind and activate the small GTPase <u>A</u>DP-<u>r</u>ibosylation factor (Arf)6 (Hofmann et al., 2007; Matsumoto et al., 2014). In the nervous system, Arf6 participates in dendrite and axon outgrowth and branching (Albertinazzi et al., 2003; Eva et al., 2012; Hernandez-Deviez et al., 2004; Miura et al., 2016), apical adhesion of neural progenitors (Arvanitis et al., 2013), and cortical neuron migration (Falace et al., 2014).

The ubiquitin-proteasome pathway plays a role in nearly every aspect of eukaryotic cell biology by targeting proteins for degradation (Glickman and Ciechanover, 2002). The large family of Cullin-RING E3 ligases (CRLs) are multi-protein complexes that nucleate around seven different Cullins. In particular, CRL5, the last evolved member of the family, assembles around Cullin 5 (Cul5) (Petroski and Deshaies 2005). On its C-terminus, Cul5 binds the RING box 2 protein (Rbx2, also known as Rnf7 or Sag), a necessary step for bringing Ub∼E2 to CRL5 and essential for full CRL5 activity (Okumura et al., 2016). To recruit substrates for ubiquitylation, CRL5 utilizes the Elongin B/C (also known as Tceb2/Tceb1) subunits to associate with up to 38 different substrate adaptor proteins, which provide substrate specificity (Okumura et al., 2016).

The core CRL5 components Cul5 and Rbx2 are ubiquitously expressed during development and in the adult central nervous system (CNS), and functionally active during development (Fairchild et al., 2018; Hino et al., 2018; Simo and Cooper, 2013). Importantly, CRL5 substrate adaptors are dynamically regulated during development, directing CRL5 activity to specific substrates at different times and areas in the CNS (Cembrowski et al., 2016). In the neocortex, CRL5 opposes projection neuron migration by downregulating the Reelin/Dab1 signaling pathway as migrating projection neurons reach the top of the cortical plate (Feng et al., 2007; Simo and Cooper, 2013; Simo et al., 2010). In the CNS, Reelin/Dab1 signaling participates in several developmental processes, including neuron migration, dendritogenesis, axogenesis, and synaptogenesis (Pujadas et al., 2010; Sekine et al., 2014; Wasser and Herz, 2017). Moreover, CRL5 also controls levels of the tyrosine kinase Fyn, which exerts important roles during development and in the adult brain (Grant et al., 1992; Knox and Jiang, 2015; Kuo et al., 2005; Simo and Cooper, 2013).

Anatomically, all mammals share a similar hippocampal sub-structure organization (Andersen, 2007). The hippocampus is divided into the dentate gyrus (DG) and *cornu ammonis* (CA). The DG includes the *fascia dentata* and the hilus, and the CA is further differentiated into CA1, CA2, and CA3. Hippocampal pyramidal neurons (PNs) are born from neural progenitors in the ventricular zone (VZ) of the cortical hem and migrate to the *stratum pyramidale* (*sp*) using radial glial-aided migration (Altman and Bayer, 1990; Khalaf-Nazzal and Francis, 2013). In comparison to the neocortex, fewer signaling pathways have been identified that control neuron migration in the hippocampus (Khalaf-Nazzal and Francis, 2013).

Here, we show that depletion of CRL5 activity disrupts localization and dendritogenesis of PNs in the hippocampus. We further demonstrate that this is a cell-autonomous phenotype and that sustained Reelin/Dab1 signaling is not sufficient to cause ectopic positioning or dendritic defects in PNs. We uncover Arl4c, Cyth-1/3, and Arf6 as novel CRL5-regulated signaling effectors and demonstrate that accumulation of Arl4c is necessary to cause defects in PN localization and dendritogenesis observed upon CRL5 inactivation. Furthermore, we show that Arf6 opposes PN over-migration and that a Cul5/Arf6 double knock-down exacerbates the over-migration phenotype. Finally, we show that Arl4c and Arf6 are necessary for PN dendrites to innervate the *stratum lacunosum moleculare* (*slm*) and, in their absence, normal apical dendrites are formed but fail to extend past the *stratum radiatum* (*sr*). Our results support a key role for CRL5 during hippocampal morphogenesis by regulating Arl4c levels and reveal a novel Arl4c/Arf6-dependent mechanism of dendritic outgrowth in PNs.

## RESULTS

### Depletion of Rbx2 disrupts projection neuron localization

To study the role of CRL5 in the developing hippocampus we crossed the previously described Rbx2 floxed mice (Rbx2 fl/fl) with Emx1-Cre mice, which expresses Cre in the telecephalon starting at embryonic stage 9.5 (E9.5) (Guo et al., 2000; Simo and Cooper, 2013). In comparison to our previous reports using Nestin-Cre to deplete Rbx2 (Rbx2cKO-Nes mice), Rbx2 fl/fl; Emx1-Cre animals (Rbx2cKO-Emx1) were normally born, thrived as control littermates, did not show the extensive hydrocephalus phenotype present in Rbx2cKO-Nes mice, and survived until adulthood. As expected, efficient and extensive depletion of Rbx2 was observed in Rbx2cKO-Emx1 samples at P1 when tested by western blotting (Figure 4C and Figure S1A). Furthermore, Rbx2cKO-Emx1 telencephalons accumulated tyrosine phosphorylated (pY)-Dab1, Dab1, and Fyn, and showed the same layering defects as previously described for Rbx2cKO-Nes pups (Figure S1) (Fairchild et al., 2018; Simo and Cooper, 2013). These results indicate that Rbx2cKO-Emx1 mice faithfully recapitulate the phenotypes observed in Rbx2cKO-Nes mice without the morbidity caused by hydrocephalus.

Depletion of Rbx2 cause a significant number of PNs, identified by staining against DKK3, to localized outside the *sp* when hippocampi from control and Rbx2cKO-Emx1 mice were analyzed at postnatal day 21-23 (P21-23) (Figure 1A and 1B) (Diep et al., 2004). Close examination identified ectopic PNs in the in the *stratum oriens* (*so*), which may be expected if migration is defective, but also in the *sr*/*slm*, suggesting that CRL5 regulates PN migration and opposes PN over-migration (Figure 1Aa’, 1Aa’’, and 1C). Most ectopic PNs were observed in the more medial part of the CA1 region with fewer ectopic PNs found in the CA3 (Figure 1A and 1C). The number of DKK3+ cells in the molecular layer and granule cell layer of the DG in the Rbx2 mutant mice was also increased (Figure 1Aa’’’,1Aa’’’’, and Figure 1C). These results indicate that Rbx2, potentially through CRL5 activity, determines the final position of PNs in the *sp* and that some PN populations, particularly those in the CA1, and DKK3+ cells in the DG, are more affected by the absence of Rbx2. Similar results were observed in Rbx2cKO-Nes hippocampi, confirming the role of Rbx2 in PN localization independently of the Cre driver used to deplete Rbx2 (Figure S2).

**Figure 1.**
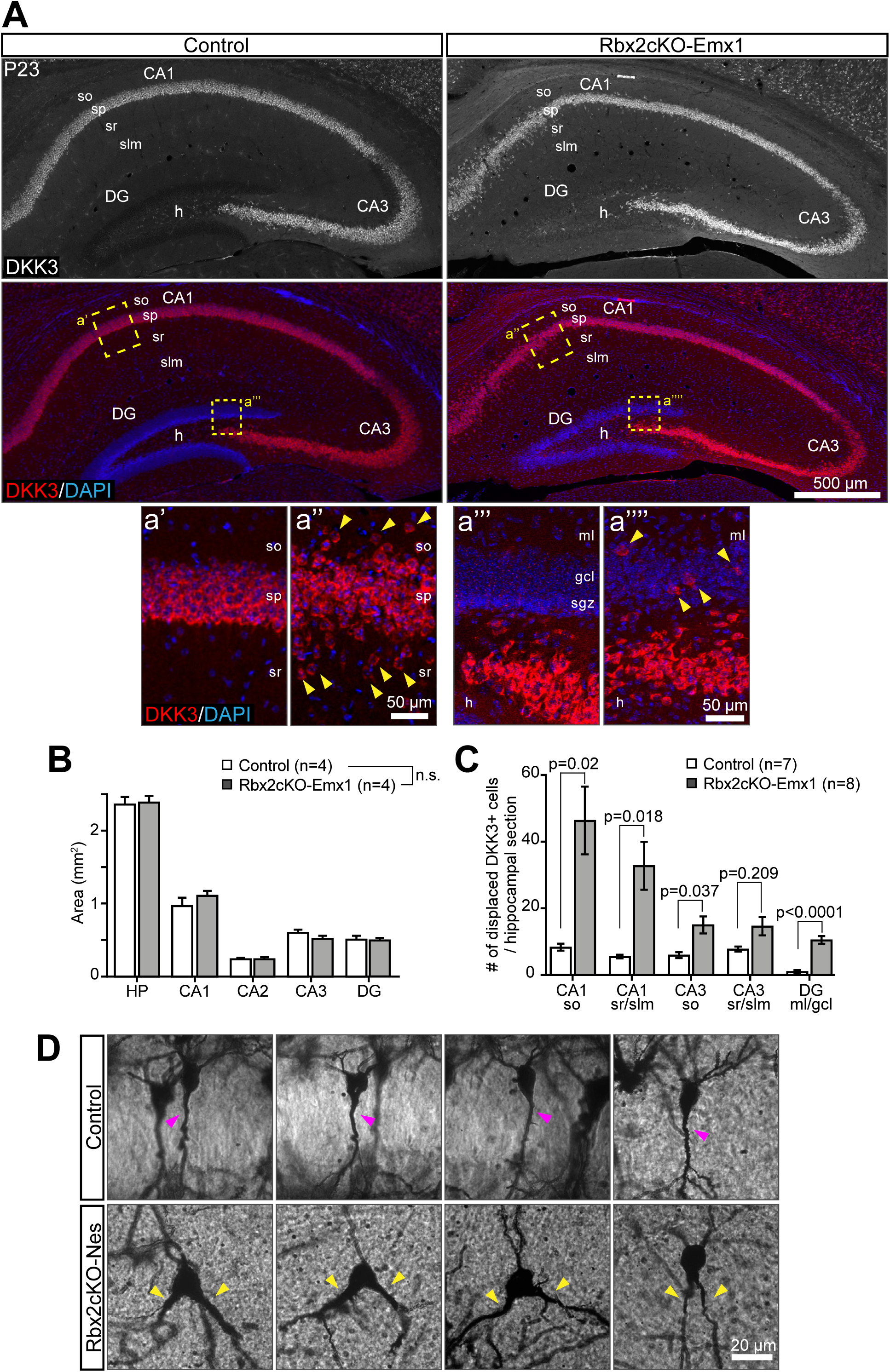
Rbx2 regulates pyramidal neuron position and apical dendrite morphology in the hippocampus. (A) DKK3+ PN misposition at P23. Stainings of coronal sections of Control (Rbx2 fl/fl) and Rbx2cKO-Emx1 dorsal hippocampus against DKK3 show displaced PNs into so, sr, and slm in Rbx2cKO-Emx1 mice. (a’-a’’’’) High-magnification images of the CA1 (a’ and a’’) and DG (a’’’ and a’’’’) showing dispersion of DKK3+ cells. Yellow arrowheads indicate displaced DKK3+ cells. (B) Quantification of displaced DKK3+ cells. Increased number of DKK3+ cells are observed in *so, sr, slm* compared to wild type littermates. 3-4 sections were quantified per brain. Mean ± SEM. Statistics, multiple t-test with Bonferroni-Dunn method adjusted p value. (C) Measurement of hippocampal areas. No differences were observed in between Rbx2 fl/fl and Rbx2cKO-Emx1 samples. Mean ± SEM. Statistics, multiple t-test with Bonferroni-Dunn method adjusted p value. (D) Abnormal multiple apical dendrites in Rbx2 mutant PNs. P15 control and Rbx2cKO-Nes P15 were stained using the Golgi silver impregnation method. “V-shaped” double apical dendrites were observed in Rbx2cKO-Nes hippocampi. Pink arrowheads indicate normal apical dendrite and yellow arrowheads indicate double apical dendrites. so: stratum oriens, sp: stratum pyramidale, sr: stratum radiatum, slm: stratum lacunosum moleculare. CA: cornu ammonis, DG: dentate gyrus, h: hilus, gcl: granule cell layer, ml: molecular layer, n.s.: not significant.

To determine whether PN morphology was affected by Rbx2 depletion, Rbx2 mutant (Rbx2cKO-Nes) and control (Rbx2 fl/fl) brains were stained with the Golgi-Cox impregnation method. In control samples, labeled PNs were found in the *sp* with a single apical dendrite perpendicular extending towards the *sr*/*slm* (Figure 1D, purple arrowhead). In comparison, several Rbx2 mutant PNs, identified by the size of their soma and location in the *sp* and *sr*, had two main (e.g., “apical”) dendrites originating at the soma and extending towards the *sr*/*slm* (Figure 1D, yellow arrowheads). These results indicate that CRL5 participates in the formation of a single apical dendrite in PNs during hippocampal development.

### CRL5 regulates pyramidal neuron localization in a cell-autonomous fashion

Our previous studies indicate that Rbx2 regulates neuron positions in the retina through cell autonomous and non-autonomous mechanisms (Fairchild et al., 2018). To test whether CRL5 regulates hippocampal development cell autonomously, we used *in utero* electroporation (IUE) to deplete CRL5 activity in individual PNs. A plasmid co-expressing a short hairpin RNA against Cul5 (shCul5) and EGFP was IUEd in CD-1 embryos at embryonic day (E) 15.5, and electroporated brains were collected at P10. In comparison to controls (GFP only), depletion of Cul5 disrupted PNs localization (Figure 2). Whereas PNs expressing GFP were mostly localized in the *sp* (91.4±1%) (Figure 2A and 2C), a significant amount of PNs expressing the shCul5/GFP plasmid were found in the *sr/slm* (73.5±0.5%) (Figure 2B and 2C). Surprisingly, few GFP+ PNs were found in the *so* when Cul5 was knocked down (KD) (0.3±0.3%) (Figure 2B and 2C), suggesting that the migration defects observed in the Rbx2cKO-Emx1 hippocampi are a combination of cell autonomous (PN over-migration) and non-cell autonomous (PN migration arrest).

**Figure 2.**
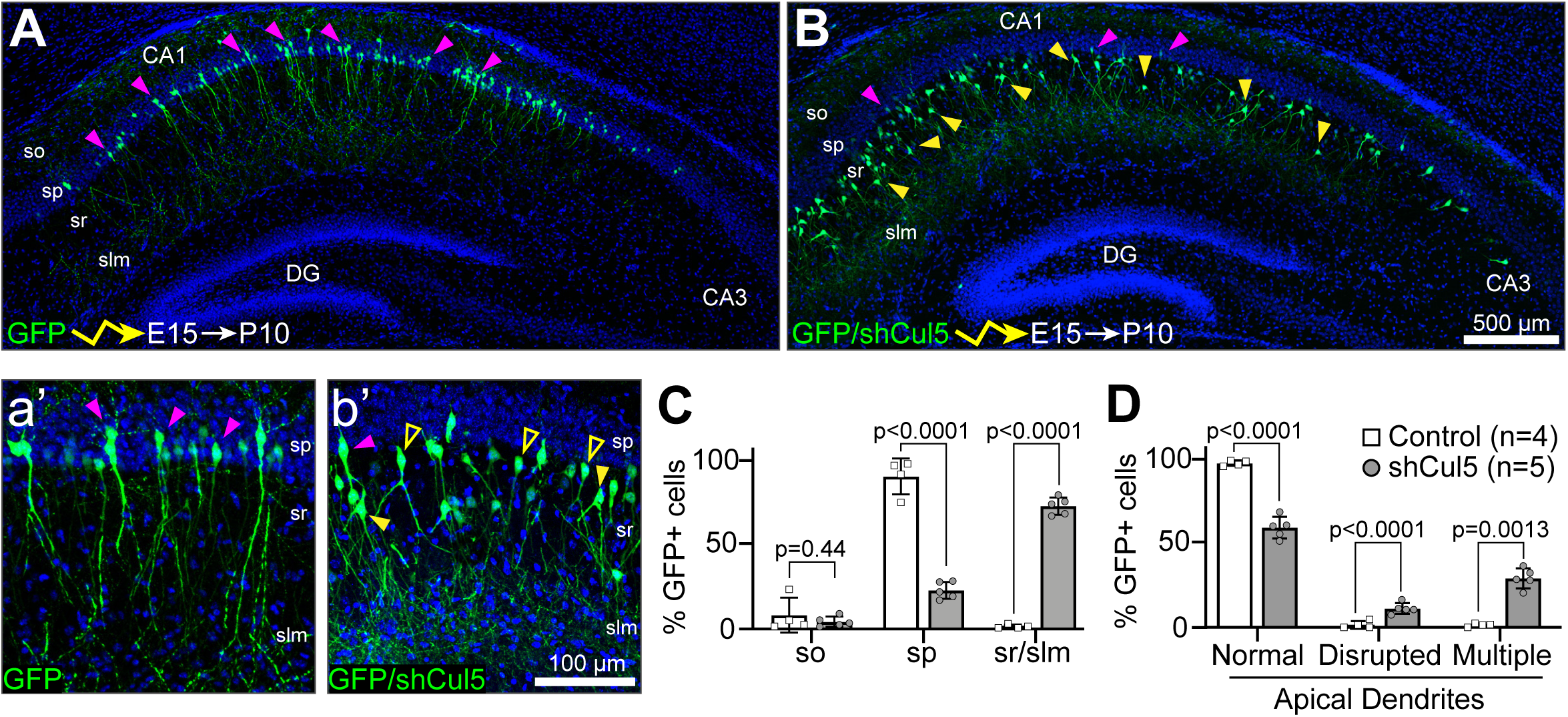
CRL5 regulates pyramidal neuron position and apical dendrite morphology in a cell-autonomous fashion. (A-B) *In utero* electroporation of GFP (A) or GFP/shCul5 (B) expressing plasmids at E15.5 embryos and samples collected at P10. shCul5 electroporated neurons over-migrated into *sr*/*slm* and had “V-shaped” multiple apical dendrites. (a’-b’) High-magnification images of A-B shows detailed PN morphology and dendritic formation. Pink arrowheads indicate PNs with correct position and apical dendrite. Yellow arrowheads indicate over-migrated PNs with abnormal apical dendrites. Yellow hollow arrowheads indicate over-migrated PNs with normal apical dendrite. (C-D) Quantification of the percentage number of displaced GFP+ cells/total GFP+ cells (C), and percentage number of GFP+ cells with normal, disrupted (single apical dendrite with abnormal orientation), and multiple (2 or more) apical dendrites in electroporated hippocampal samples. At least 3 sections were quantified per electroporated brain. Mean ± SD. Statistics, multiple t-test with Bonferroni-Dunn method adjusted p value.

Next, we analyzed the apical dendrites in the electroporated PNs. As expected, the majority of GFP+ PNs in control electroporations showed a normal, single apical dendrite extending radially towards the *sr* and *slm* (Figure 2a’ and 2D). Depletion of Cul5 disrupted dendrite formation in more than 40% of the PNs analyzed (Figure 2b’ and 2D). Two main defects were observed: 1) a disrupted apical dendrite, where a single apical dendrite is not radially oriented towards the *sr*/*slm*; and 2) PNs with two or more “apical” dendrites (i.e. Multiple apical dendrites) (Figure 2b’ and D). Interestingly, in the majority of cases where PNs had two “apical” dendrites, these dendrites formed a “V-shape” extending to the *sr* and the *slm*. These results are consistent with those observed in Rbx2cKO-Nes mice (Figure 1D) and indicate that CRL5 participates in the formation of a single, radially oriented apical dendrite in hippocampal PNs.

### Sustained Reelin/Dab1 signaling does not cause PN over-migration or dendritic defects

CRL5 opposes Reelin/Dab1 signaling by targeting active pY-Dab1 for degradation (Feng et al., 2007). Reelin/Dab1 signaling is believed to regulate PN migration and dendrite outgrowth (Jiang et al., 2016; Niu et al., 2004). Thus, we hypothesized that sustained Reelin/Dab1 signaling might be responsible for the phenotypes observed in the Rbx2cKO mutant hippocampus. First, we confirmed that pY-Dab1/Dab1 accumulated in the Rbx2cKO-Emx1 hippocampus (Figure 4C) and in shCul5-electroporated neurons (Figure S3A). To test whether sustained Reelin/Dab1 signaling is sufficient to promote PN over-migration, we used the SOCS7 knock out mouse (Fairchild et al., 2018). SOCS7 is a CRL5 substrate adaptor that recruits and targets pY-Dab1 for degradation in the developing telencephalon (Simo and Cooper, 2013). Importantly, depletion of SOCS7 promoted an accumulation of Dab1 in the hippocampus at similar levels as in Rbx2 mutant animals, but without affecting other CRL5-dependent substrates (Figure 3A, 3B and S3B). Next, we analyzed the distribution of DKK3+ PNs in the SOCS7 mutant hippocampus. No differences were observed in PN position between control (SOCS7 +/-) and SOCS7 mutant hippocampi (Figure 3C and 3D). Furthermore, we used IUE to express GFP in control and SOCS7 mutant PNs. The majority of GFP+ cells were found in the *sp* both in control and SOCS7 mutant hippocampi (Figure 3E). Importantly, in both cases, PNs present a single, apical dendrite that extends radially to the *sr*/*slm*. These results indicate that sustained signaling from Reelin/Dab1 or from other SOCS7 targets are not sufficient to promote the PN over-migration or dendritic defects observed in the Rbx2 mutant hippocampus.

**Figure 3.**
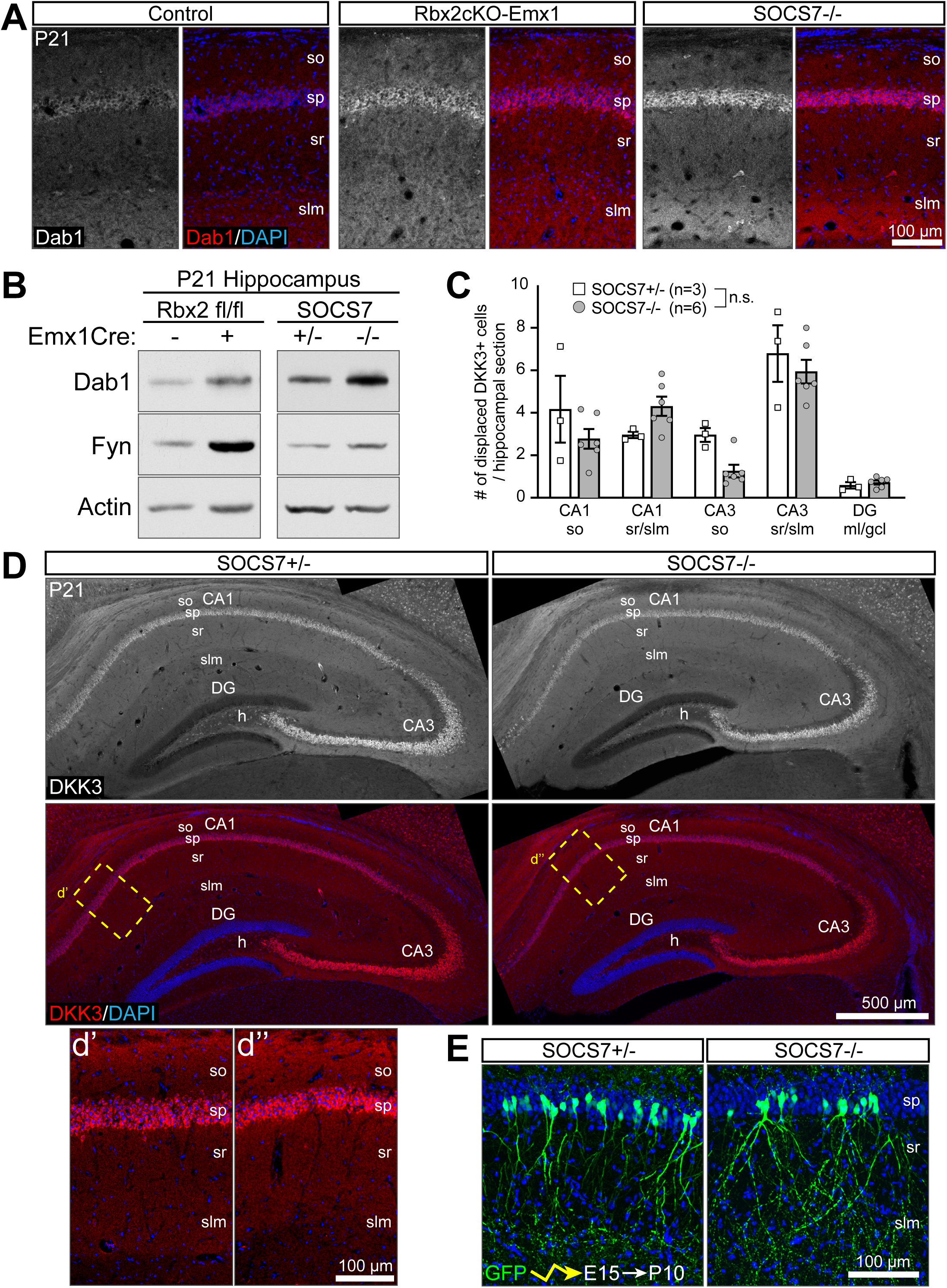
Dab1 accumulation is not sufficient to disrupt pyramidal neuron positioning or apical dendrite morphology. (A) Dab1 accumulated in both Rbx2cKO-Emx1 and SOCS7-/- hippocampi in comparison to control (Rbx2 fl/fl). Coronal sections of P21 hippocampi were stained with anti-Dab1 and counterstained with DAPI. (B) Representative images of western blotting experiments indicating that Dab1 accumulated in both Rbx2cKO-Emx1 and SOCS7-/- hippocampal samples, but Fyn only accumulated in Rbx2cKO-Emx1 P21 hippocampal lysates. (C) Quantification of displaced DKK3+ cells. No difference was observed between genotypes. At least 3 sections quantified per brain. Mean ± SD. Statistics, multiple t-test with Bonferroni-Dunn method adjusted p value. (D) Coronal sections of P21 SOCS7+/- and SOCS7-/- hippocampi were stained with an anti-DKK3 antibody and counterstained with DAPI. (d’-d’’) High-magnification images of the CA1 indicating that normal PN localization in SOCS7-/- hippocampi. (E) *In utero* electroporation of GFP in SOCS7+/- and SOCS7-/- embryos at E15.5 and collected at P10. Electroporated GFP+ PNs show normal positioning and apical dendrites.

**Figure 4.**
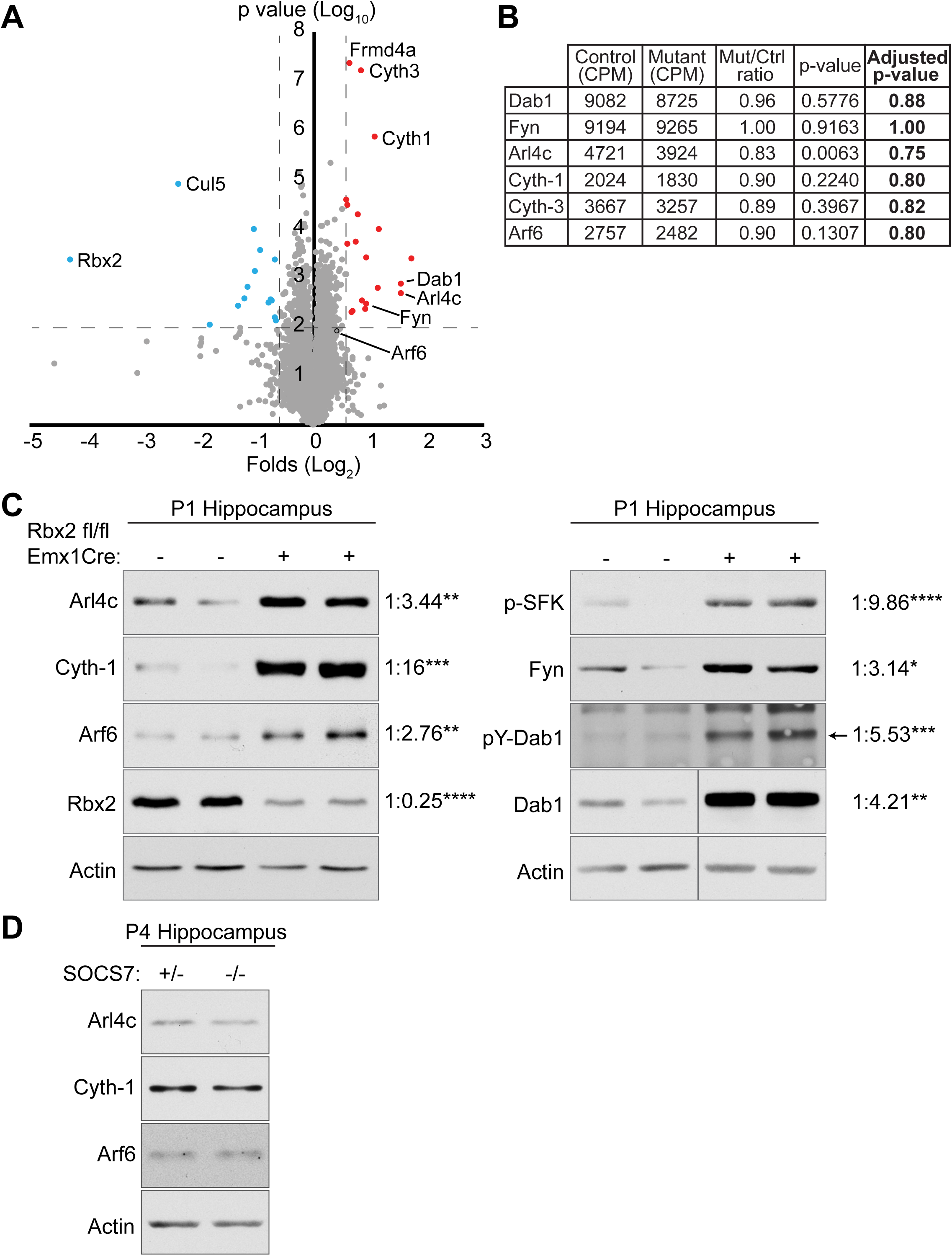
Arl4c, Cytohesin-1, and Arf6 accumulate in the Rbx2 mutant but not in SOCS7-/- hippocampi. (A) Tandem Mass-Tag mass spectrometry (TMT-MS) of Rbx2cKO-Nes (n=3) and Rbx2 fl/fl (n=4) telencephalons (P0) plotted as a ratio of Rbx2cKO-Nes/control. X axis represents the logarithmic fold ration of Rbx2cKO-Nes/control per protein identified. Y axis represents the logarithmic p value of the t test per protein identified. Proteins whose differentially expression in comparison to control (Rbx2 fl/fl) have a p<0.01 and accumulated above 1.5 folds (red dots) or reduced lower than 0.666 folds (blue dots) in comparison to control were initially considered for further studies. As expected, Rbx2 mutant samples showed lower levels of Rbx2 and Cul5 and higher levels of Dab1 in comparison to control (Simo and Cooper, 2013). (B) RNA sequencing analysis comparing Rbx2cKO-Nes (Mutant; n=3) and Rbx2 fl/fl (Control; n=3) telencephalon at P0 was performed to determine transcriptionally differences between genotypes. Table shows the sequencing results of genes encoding for the proteins identified in (A). P values were calculated using t tests and Adjusted p-values were calculated using the False Discovery Rate method. No significant transcriptomic differences were observed between genotypes. CPM: counts per million. (C) Western blot analyses validating TMT-MS data in P1 Rbx2cKO-Emx1 and control hippocampi. Arrow indicates pY-Dab1 band in anti-p-Tyr blot. Numbers indicate ratio of proteins in Rbx2cKO-Emx1 (n=3) vs Rbx2 fl/fl (n=5). Statistics, Student’s t-test. *: p<0.05, **: p<0.01, ***: p<0.001, ****: p<0.0001. (D) Arl4c, Cyth-1, and Arf6 did not accumulate in P4 SOCS7 -/- (n= 3) in comparison to controls (SOCS7 +/-, n= 4) hippocampus lysates when analyzed by western blotting. Statistics, Student’s t-test.

### CRL5 regulates levels of Arl4c, Cytohesin-1, and Arf6 in the developing telencephalon

To determine which proteins and signaling pathways are responsible for the phenotypes observed in the Rbx2 mutant hippocampus, we performed quantitative mass spectrometry (MS) using the tandem mass tag isobaric-labelling comparing control (Rbx2 fl/fl, n=4) and Rbx2 mutant (Rbx2cKO-Nes, n=3) P1 telencephalons (Thompson et al., 2003). 8088 unique proteins with two or more peptides identified per protein were considered for further analysis (Figure 4A and Supplementary Table 1). Proteins accumulated or decreased 1.5-fold in comparison to control samples and with Student t-test p<0.01 were primarily considered as regulated by CRL5. As previously shown, Rbx2 and Cul5 were downregulated and the bona fide CRL5 targets Dab1 and Fyn accumulated in the Rbx2 mutant samples (Simo and Cooper, 2013). The same results were observed in hippocampal samples from Rbx2cKO-Emx1 mice when levels of Rbx2, pY-Dab1/Dab1, and p-SFK/Fyn were analyzed by western blotting (Figure 4C). These results further confirm that, at the molecular level, Rbx2, potentially through CRL5 activity, regulates Dab1 and Fyn levels independently of the Cre driver mouse used to knock out *rbx2*. We observed the accumulation of 20 proteins in the Rbx2 telencephalons, including Arl4c, Cyth-1, and Cyth-3 and Arf6 (Figure 4A). Importantly, we confirmed by western blotting that Arl4c, Cyth-1, and Arf6 significantly accumulated in the hippocampus of Rbx2cKO-Emx1 mice at perinatal stages (Figure 4C), suggesting that CRL5 regulates levels of Arl4c and Arl4c signaling effectors in the hippocampus. However, these results cannot determine whether CRL5-dependent regulation is a direct (e.g., recruiting, poly-ubiquitylating, and targeting effectors for degradation) or indirect (e.g., affecting levels of gene transcription) phenomenon. To test whether transcription was affected in Rbx2 mutant mice, we compared the transcriptome of control and Rbx2cKO-Nes telencephalons (P1) by RNA sequencing. Absence of Rbx2 did not significantly alter the expression profile of any detected gene, including Dab1, Fyn, Arl4c, Cyth-1/3, or Arf6 (Figure 4B). These results indicate that CRL5-dependent regulation of Arl4c, Cyth-1/3, and Arf6 is post-transcriptional.

### Arl4c is dynamically expressed during hippocampal development

To determine the role of Arl4c in the developing murine hippocampus, we first determined its ontogenic expression. Arl4c was expressed in the hippocampus with maximal expression between P5 and P9, which correlates with PN apical dendrite outgrowth, and minimal expression at embryonic and juvenile stages (Figure 5A and 5B). These changes in protein expression are driven by post-transcriptional mechanisms as no significant differences in Arl4c mRNA levels were detected during postnatal development and in the adult hippocampus (Figure 4B and 5C). Importantly, absence of Rbx2 promotes Arl4c accumulation at any stage of development, suggesting that CRL5 is constantly regulating Arl4c levels (Figure 5D). Immunofluorescence against Arl4c in a P8 mouse brain indicated that Arl4c is ubiquitously expressed in the brain (Figure S4). Expression of Arl4c was detected in the cytoplasm of PNs and granule cells of the dentate gyrus of control and Rbx2 mutant hippocampi, albeit at a higher level in the latter (Figure 5E and S4).

**Figure 5.**
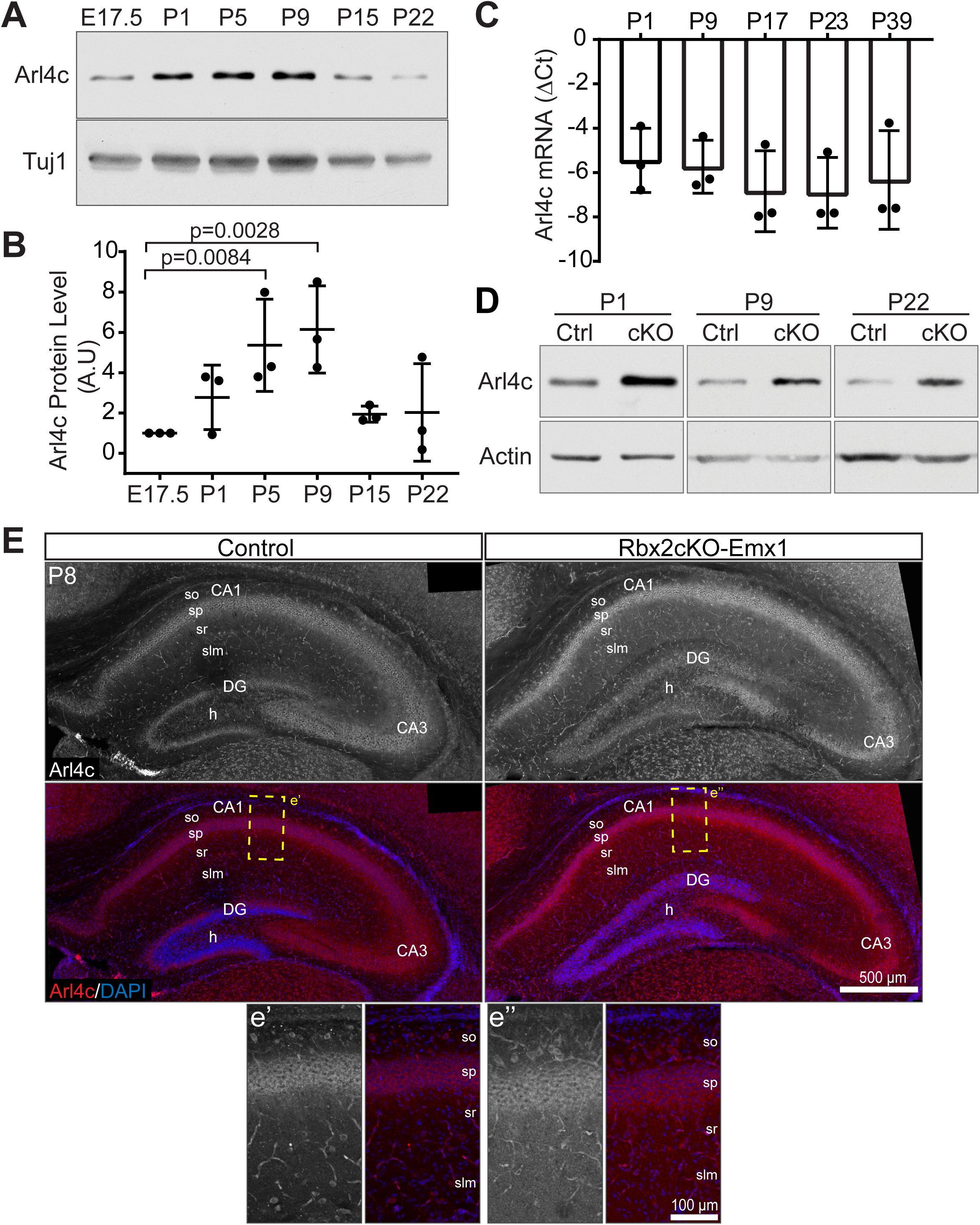
Arl4c is dynamically expressed during hippocampal development. (A) Ontogenic analysis of Arl4c expression in the hippocampus during development by western blotting. (B) Quantification of Arl4c western blots in (A) and normalized to Arl4c expression at E17.5. Mean ± SD. Statistics, one-way ANOVA. (C) No changes were observed in *arl4c* transcript levels between ages P1 to P39 by qPCR analysis. Cycle different between Arl4c and Actin from three independent experiments are plotted. Mean ± SD. Statistics, one-way ANOVA. (D) Arl4c levels from hippocampal lysates of Rbx2 fl/fl (Ctrl) and Rbx2cKO-Emx1 (cKO) were compared at several ages by western blotting. Arl4c accumulated in Rbx2cKO-Emx1 samples at all ages analyzed. (E) Coronal sections of Control (Rbx2 fl/fl) and Rbx2cKO-Emx1 at P8 were stained with anti-Arl4c antibody and counterstained with DAPI. Arl4c is highly expressed in PNs of the CAs. (e’-e’’) High-magnification images of the CA1 regions indicated in (E).

In hippocampal PNs, Arl4c was detected in puncta, mostly in the cytoplasm but also in the nucleus, and diffused at the plasma membrane (Figure S5A). To determine whether Arl4c localization was changing in response to Rbx2 depletion, we cultured E17.5 hippocampal neurons from control and Rbx2cKO-Emx1 embryos and analyzed Arl4c expression at four days in vitro (DIV). In culture neurons from control hippocampi, Arl4c was detected in puncta in the cytoplasm, both in the soma and neurites, as well as at the plasma membrane (Figure S5B). Rbx2 mutant neurons showed a similar Arl4c distribution as in the controls but with an increased Arl4c signal both at the plasma membrane and in puncta (Figure S5C). Interestingly, depletion of Rbx2 significantly increased the total number of Arl4c+ puncta in the cell (Figure S5D). These results confirm that depletion of Rbx2 promotes Arl4c accumulation at the single-cell level, that Arl4c is distributed both in puncta and at the plasma membrane in hippocampal neurons, and suggest that neurons oppose the increased levels of Arl4c by storing it in intracellular vesicles.

### Arl4c participates in dendritic development *in vitro*

Given that Arl4c expression correlates with dendrite formation and maturation in hippocampal pyramidal neurons and that Arl4c regulates cytoskeleton dynamics in epithelial cell lines (Chiang et al., 2017; Nakahira and Yuasa, 2005), we assessed whether Arl4c participates in PN dendritogenesis. E17.5 hippocampal primary neurons were co-transfected with a hARL4C expression plasmid or a shArl4c expressing plasmid along with an EGFP expression plasmid (Figure 6A and S6A). We fixed the cells after two days and measured neurite complexity by Sholl analysis. In comparison to controls, over-expression of hARL4C significantly reduced neurite complexity (Figure 6A and 6B). On the contrary, depletion of endogenous Arl4c increased neurite complexity (Figure 6A and 6B). Interestingly, when the distance of the longest neurite was measured, which usually corresponds to the developing axon, no differences were observed between conditions (Figure 6C), nor was the complexity of the longest neurite affected in any condition (Figure 6B, >60µm). These results suggest that Arl4c participates in the formation of the dendritic tree in hippocampal neurons without affecting the growth or complexity of the prospective axon.

**Figure 6.**
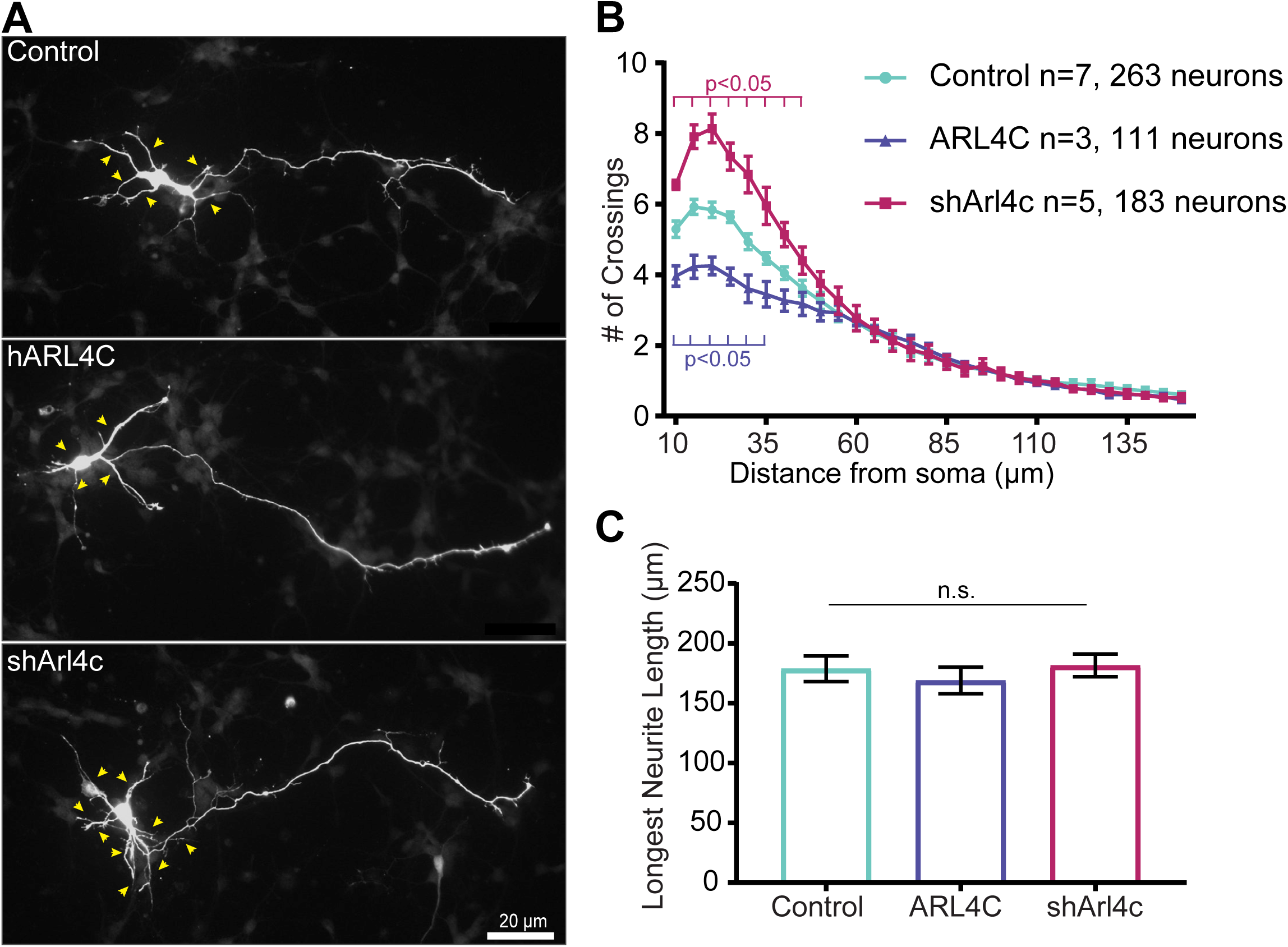
Arl4c participates in dendritic development of cultured hippocampal neurons. (A-B) Arl4c expression negative regulate dendritic complexity of hippocampal neurons. E16 hippocampal primary neurons were obtained and transfected in suspension with GFP (Control) or human ARL4C /GFP (hARL4C) expression plasmids, or a shRNA against Arl4c/GFP (shArl4c) plasmids. Representative images (A) and Sholl analysis (B) from independent experiments are shown. 20-50 neurons were quantified per experiment. ARL4C over-expression decreased and Arl4c knockdown increased dendritic complexity within 60 μm of soma. Yellow arrowheads indicate neurites. Mean ± SEM. Statistics, two-way ANOVA. (C) Length of the longest neurites were measured in each condition and no statistical differences were observe between conditions (n=3). Number of neurons quantified: 125- Control; 119-ARL4C; 168-shArl4c. Mean ± SEM. Statistics, one-way ANOVA.

### Arl4c and Arf6 regulate pyramidal neuron migration and dendrite formation during hippocampal development

Finally, we tested the possibility that accumulation of Arl4c and/or Arf6 in PNs was responsible for the migration and dendritic phenotypes observed upon CRL5 inactivation (Figure 7A and 7B). Using IUE, we knocked down Cul5 together with Arl4c or Arf6 from hippocampal PNs (Figure 7C and 7D, and Figure S6). Depletion of Arl4c completely rescued the over-migration phenotype caused by Cul5 KD and the majority of PNs were found in the *sp* with a single apical dendrite towards the *sr* (Figure 7C and 7E). However, the dendritic tree of Cul5/Arl4c-KD PNs failed to extend into the *slm*. Apical dendrites from Cul5/Arl4c-KD PNs emerged normally but an obvious increase in dendritic arborization occurred at the lower end of the *sr* and did not innervate the *slm* (Figure 7C).

**Figure 7.**
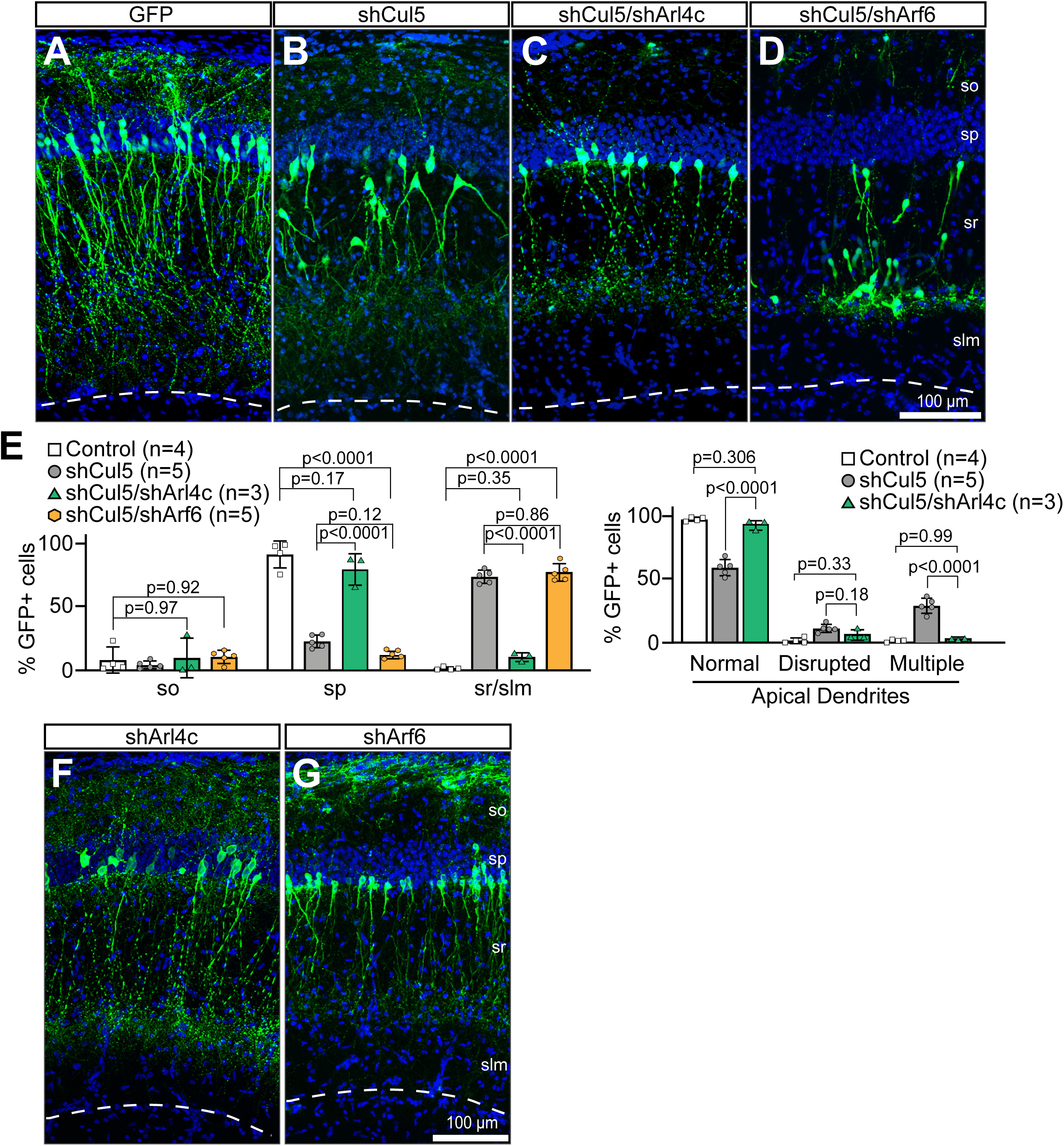
Arl4c and Arf6 have opposite effects on CRL5-dependent PN over-migration but same effects on PN dendrite extension into the stratum lacunosum moleculare. (A-D) Representative images of hippocampus electroporated at E15 and collected at P10 showing (A) control (GFP), (B) Cul5 KD (shCul5), (C) double Cul5/Arl4c KD (shCul5/shArl4c), and (D) double Cul5/Arf6 KD (shCul5/shArf6) transfected PNs. Whereas Arl4c KD rescue the PN over-migration phenotype caused by Cul5 KD, Arf6 KD exacerbated it. Both Cul5/Arl4c- or Cul5/Arf6-KD PNs fail to extend their apical dendrites into the *slm* in comparison to control or Cul5-KD PNs. The striped lines represent the hippocampal fissure. (E) Quantification of GFP+ cells in each strata (left) and apical dendrite morphology (right). Statistics, multiple Student t-test with Bonferroni-Dunn method adjusted p value. (F-G) Single (F) Arl4c (shArl4c) or (G) Arf6 (shArf6) KD constructs were co-transfected with GFP by in utero electroporation in E15 hippocampi and brains were collected at P10. In both conditions, PNs migrated normally, had a single radially-oriented apical dendrite, but fail to extend their dendrites into the *slm*. The striped lines represent the hippocampal fissure.

Knocking down Arf6 did not rescue the PN over-migration phenotype caused by Cul5 KD, but exacerbated PN over-migration with more PNs located the at lower end of the *sr* (Figure 7D and 7E). These results support the hypothesis, previously established in the cortex, that active Arf6 inhibits neuron migration (Hara et al., 2016). Despite the enhancement in PN displacement, no Cul5/Arf6-KD PN was found in the *slm* and in several instances electroporated PNs were found arrested at the boundary between *sr* and *slm*. Due to the large clustering of Cul5/Arf6-KD PNs at the *sr*/*slm* boundary, we were unable to precisely determine the dendrite morphology of these neurons. However, we did not observe any dendrite penetrating into the *slm* (Figure 7D). These data suggest that CRL5 participates in PN localization and apical dendrite formation by regulating Arl4c and Arf6 levels.

Given that Cul5-KD PNs did extend dendrites into the *slm*, we hypothesized that the failure to do so in Cul5/Arl4c- and Cul5/Arf6-KD PNs would be independent of CRL5 activity but dependent on Arl4c and Arf6 activity. To test this hypothesis, we IUEd shArl4c and shArf6 individually in the hippocampus and determined the ability of electroporated PNs to innervate the *slm*. Both Arl4c and Arf6-KD PNs were located in the *sp* and extended a single apical dendrite towards the *sr* (Figure 7F and 7G). Importantly, depletion of Arl4c or Arf6 blocked, for the most part, the ability of PNs to extend their dendrites into the *slm*, confirming that Arl4c and Arf6 are novel signaling effectors involved in PN dendritic outgrowth into the *slm*.

## Discussion

Our results show that CRL5 is required for PN migration and apical dendrite formation. Rbx2 mutant PNs, principally those from the CA1 region, are ectopically displaced into the *so* and *sr*/*slm*. Over-migration into the *sr*/*slm* is a cell-autonomous effect of CRL5 inactivation, but cell autonomous Cul5 KD does not cause under-migration into the *so*. These results indicate that both cell and non-cell autonomous phenotypes shape the Rbx2 mutant hippocampus. Moreover, CRL5 inactivation disrupts the formation of normal, single, radially oriented PN apical dendrites and promotes the formation of multiple or non-radially oriented apical dendrites. We uncovered Arl4c as a novel signaling effector present in PNs, dynamically expressed during brain development, and important for dendritic arborization in hippocampal neurons. Furthermore, RNA sequencing and quantitative proteomic assays indicate that CRL5 regulates levels of Arl4c, Cyth-1, and Arf6 post-transcriptionally, and that accumulation of Arl4c is necessary for the phenotypes observed in CRL5-depleted PNs. Finally, we show that Arl4c and Arf6 participate in the outgrowth of PN apical dendrites into the *slm*.

### CRL5 regulation of Arl4c signaling

Our quantitative proteomic analysis indicates that CRL5 is a crucial regulator of the brain proteome during development in a potential direct (proteins accumulated in absence of Rbx2) and indirect fashion (proteins loss). Among these, we identified several proteins accumulated in absence of Rbx2, including Arl4c, Cyth-1/3, Arf6, Frmd4a, and Fyn. Several examples of protein-protein interactions as well as functional interconnection have been demonstrated for these signaling effectors. Arl4c recruits all members of the cytohesin family to the plasma membrane by binding with the pleckstrin homology domain present on cytohesins. At the plasma membrane, cytohesins can recruit and activate Arf6 by promoting GDP/GTP exchange through the Sec7 domain (Hofmann et al., 2007; Ratcliffe et al., 2019). Importantly, in Schwann cells, Fyn regulates Cyth-1 GEF activity towards Arf6 by phosphorylating its Tyr^382^, a mechanism that may be present in other cytohesins (Yamauchi et al., 2012). Finally, Cyth-1 interacts with Frmd4a to activate Arf6 and regulate cell polarity, actin cytoskeleton, and membrane trafficking in epithelial and neuronal cells (Ikenouchi and Umeda, 2010; Yan et al., 2016). The functional interconnection among these proteins and the fact that, in absence of CRL5 activity, all of them accumulated in the telencephalon indicates that CRL5 is a crucial regulator of Arl4c and Arl4c signaling.

Different to other Arls and Arfs, Arl4c has a spontaneous GTP binding capability that enables Arl4c to be constantly active (Chiang et al., 2017; Matsumoto et al., 2017). In epithelial cells, Arl4c localizes preferentially at the plasma membrane, where it interacts with cythohesins, or in the nucleus (Hofmann et al., 2007; Matsumoto et al., 2017). Interestingly, our results indicate that Arl4c localizes at the plasma membrane and in punctum in the cytoplasm whereas little Arl4c signal is detected in the nucleus. Given that no GAPs or GEFs has been described for Arl4c (Sztul et al., 2019), our results that Rbx2 mutant hippocampal neurons have an increased number of Arl4c-containing punctum suggests that Arl4c activity may be regulated by CRL5-dependent turnover or by sequestering Arl4c in cytoplasmic vesicles and away from the plasma membrane.

Arl4c promotes epithelial cell migration (Chiang et al., 2017; Matsumoto et al., 2014). Accordingly, our IUE experiments show that reducing Arl4c levels in Cul5-depleted PNs rescued PN over-migration. However, during normal PN migration endogenous levels of Arl4c are relatively low and depletion of Arl4c do not affect PN positioning in the *sp*. In the neocortex, Arf6 participates in the multipolar-to-bipolar transition of migrating cortical neurons and Arf6 inactivation (i.e. Arf6-GDP) is necessary for cortical neuron migration (Falace et al., 2014; Hara et al., 2016). Similarly, inactivation of Arf6 is necessary to maintain cortical neural progenitors’ polarity, which affects neurogenesis and neuron migration in the cortex (Arvanitis et al., 2013). Our results suggest a model where accumulation of Arl4c recruits Cyth-1/3 to the plasma membrane, promoting cell-migration, and at the same time activates Arf6, which partially opposes over-migration. Then, depletion of Arf6 enhances PN over-migration caused by Cul5 KD. Finally, depletion of Arf6 alone does not affect normal hippocampal PN migration, indicating different roles for Arf6 during hippocampal and cortical neuron migration and underscoring that the molecular mechanism controlling projection neurons in the cortex and hippocampus are not equal.

### CRL5 controls pyramidal neuron migration and apical dendrite formation

Our results indicate that inactivation of CRL5 differentially affects different populations of PNs in the hippocampus. Similar results were observed in Rbx2 mutant retinas where some rod bipolar cells and cone photoreceptors were displaced whereas others remain in their intended layers (Fairchild et al., 2018). Also, in the Rbx2 mutant neocortex, all Ctip2+ neurons are significantly more disorganized than Tbr1+ or Cux1+ neurons (Simo and Cooper, 2013). These results suggest that some neurons are more sensitive to the loss of CRL5, eventually defining a new subpopulation among sibling neurons. Given that the expression of Cul5 and Rbx2 is homogenous within these neuronal populations (Hino et al., 2018), the molecular differences that sensitize CA1 PNs, and other neuronal types, remain to be elucidated.

Besides its role on neuron position, CRL5 participates in PN apical dendrite formation. The “v-shape” dendritic phenotype described in Rbx2 mutant and Cul5 KD PNs demonstrates the cell-autonomous role of CRL5 in PN apical dendrite formation. V-shape dendrites have been previously described in granule cells of the mouse hippocampus in response to stress, inflammation, or in neurodegenerative models, including models of Alzheimer’s Disease (AD) and Frontotemporal dementia, as well as in the brains of AD patients (Llorens-Martin et al., 2013; Llorens-Martin et al., 2015; Terreros-Roncal et al., 2019). Considering that Cul5 acts cell-autonomously and that reducing Arl4c levels, in IUE experiments, rescues this phenotype, it is unlikely that stress or inflammation may be responsible for the phenotypes observed upon CRL5 inactivation. Tau secretion has also been shown to promote “v-shape” dendrites in hippocampal granule cells (Bolos et al., 2017). Interestingly, Cyth-1 and Frmd4a modulates Tau secretion (Yan et al., 2016) and Cyth-1, Frmd4a, and Tau accumulate in the Rbx2 mutant telencephalons (Supplementary Table 1), suggesting that, in the absence of CRL5, Tau secretion may be enhanced, promoting the “v-shape” dendritic phenotype.

Our work also uncovered Arl4c as a novel, functional small GTPase in the nervous system. Moreover, its dynamic protein expression, but stable mRNA expression, in the hippocampus over time indicates a tight post-transcriptional regulation, most likely by CRL5-dependent turnover. Given the low levels of Arl4c during the period of PN migration (Khalaf-Nazzal and Francis, 2013) and Arl4c KD experiments (Figure 7F), our results convey that Arl4c does not participate in PN migration and only its strong accumulation in absence of CRL5 activity may be responsible for the migration defects observed in PNs. However, our data indicate that Arl4c participates in PN dendritogenesis *in vitro* and *in vivo. In vivo*, Arl4c and Arf6 regulate PN dendrite outgrowth into the *slm*. Interestingly, endogenous Arl4c expression peaks during PN dendritogenesis at early postnatal stages. We hypothesize that increase levels of Arl4c are necessary to activate Arl4c signaling, triggering dendritic outgrotwth into the *slm*.

## Materials and Methods

### Animals

All animals were used with approval from the University of California, Davis Institutional Animal Care and Use Committees and housed and cared for in accordance with the guidelines provided by he National Institutes of Health. *rbx2* floxed (Rbx2 fl/fl), *rbx2 fl/fl; Nestin-Cre/+* (Rbx2cKO-Nes) mice were obtained as previously described (Simo and Cooper 2013). To generate *rbx2 fl/fl; Emx1-Cre/+* (Rbx2cKO-Emx1) mice, *rbx2 fl/fl* were crossed with *Emx1-Cre/+* mice (Guo et al., 2000) and heterozygous mice for both alleles were crossed with *rbx2 fl/fl.* Rbx2cKO-Emx1 mice were viable, fertile and survive until adulthood without any evidence of hydrocephalus. *socs7* heterozygous and knock-out mice were obtained as previously described (Fairchild et al 2018). CD-1 mice were used for IUE experiments and Sholl analysis (Charles River). Females were mated and the morning a vaginal plug was observed was considered embryonic day 0.5 (E0.5). In utero microinjection and electroporation were performed as previously described (Simo and Cooper, 2013).

### Constructs

The pSG5-hARL4C wild-type construct was a generous gift from Dr. Fang-Jen Lee (National Taiwan University, Taiwan) (Chiang et al., 2017). pcDNA3-HA-Arf6 was a gift from Thomas Roberts (Addgene plasmid # 10835, (Furman et al., 2002)). For our experiments, both hARL4C and HA-Arf6 were subclone into a pCAG vector (Simo and Cooper, 2013). Short-hairpin RNAs were clone into pSUPER (Oligoengine) according to manufacturer recommendations (shArl4c 5’- AGTTCACACAGAAACCCTGGG-3’; shArf6 5’-AGCTGCACCGCATTATCAA-3’ (Kim et al., 2015)) or into pMX-puro (shCul5 5’-GCTGCAGACTGAATTAGTAG-3’ (Simo and Cooper, 2013)).

### Histology and Immunohistochemistry

Two-week old and older mice were anaesthetized and transcardially perfused with phosphate buffer saline (PBS) followed by 3.7% formalin/PBS using a peristaltic pump. Perfused fixed brains were collected and post-fixed at 4°C overnight in the same solution. Tissues were cryoprotected with 30% sucrose/PBS solution. Next, brains were embedded in Optimum Cutting Temperature (OCT) compound (Tissue-Tek, Torrance, CA) and quickly frozen using dry-ice. OCT embedded brain blocks were cryo-sectioned on a coronal plane (30 µm). Immunostainings were performed in free-floating sections with agitation. Sections were blocked with PBS, 0.5% Triton X-100 and 5% milk or 10% normal donkey serum for 1h at room temperature. Blocking solution but reducing Triton X-100 concentration to 0.3% was used for primary antibody incubation (overnight, 4°C). The following primary antibodies were used for immunohistochemistry: anti-Ctip2 (1:400, Abcam #ab18465), anti-Cux1 (1:50, Santa Cruz Antibodies #sc-13024, Discontinued), anti-Dkk3 (1:200, SinoBiological #50247-RP02), anti-Dab1 (1:100, Sigma-Aldrich #HPA052033), anti-Arl4c (1:100, Proteintech #10202-1), anti-MAP2 (1:200, Abcam #5392), and anti-GFP (1:200, Life Technologies #A11122). After primary antibody incubation, free-floating sections were washed three times in PBS/0.1% Triton X-100 (10 min each). Species-specific AlexaFluor 488-, 568-, and/or 647-conjugated IgG (1:200; Invitrogen) were used in blocking solution but reducing Triton X-100 concentration to 0.3% (90 minutes, room temperature). 4’,6-diamidino-2-phenylindole (DAPI) (Sigma-Aldrich) was used for nuclear staining. Images were taken in a Fluoview FV3000 confocal microscope (Olympus) or Axio Imager.M2 with Apotome.2 microscope system. All images were assembled using Adobe Photoshop and Illustrator.

### Golgi-Cox Stainings

Golgi-Cox stainings were performed on freshly dissected P15 Rbx2 fl/fl and Rbx2cKO-Nes brains using FD Rapid GolgiStain Kit according to the manufacturer protocol (FD NeuroTechnologies #PK401A).

### Primary cultures

E17.5 hippocampi were dissected from CD-1, Rbx2fl/fl, or Rbx2cKO-Emx1 embryos and neurons were dissociated using papain (Worthington Papain Dissociation Systems cat# LK003150). Tissue were incubated in papain for 10 minutes at 37°C with occasional agitation and triturated to further dissociate cells. Cells were pelleted and resuspended in ovomucoid protease inhibitor containing solution following the manufacturer protocol and 0.5M cells were plated on Poly-D-Lysine (0.05 μg/μl) coated glass coverslips. Neurons were transfected in suspension immediately following dissociation using Lipofectamine 2000 (ThermoFisher #11668019) following manufacturer protocol. Neurons were cultured in Neurobasal medium containing 1x P/S, 1x GlutaMAX, 30mM D-glucose, and B27 supplement. Media was changed every 2 days and neurons were fixed at indicated time points using 3.7% formalin/PBS for 15 minutes at 4°C.

### Sholl Analyses/Neurite Measurements

Hippocampal primary neurons were transfected in suspension on the day of isolation with 0.2µg of pCAG-GFP, 0.2 µg of pCAG-GFP and 1µg pCAG-hARL4C, or 0.2 µg of pCAG-GFP and 1µg pSUPER-shArl4c as indicated. Neurons were fixed at 2DIV and stained against GFP. GFP+ neurons were imaged in a Zeiss epifluoresence microscope. Neurite complexity in each condition was measured by placing a grid of concentric circles, with 5 µm radius increment, around the neuron cell body and counting the number of neurites crossing each circle. Longest neurite length was measured using ImageJ plugin Simple Neurite Tracer.

### RNA sequencing

RNA from Rbx2 fl/fl (n=3) and Rbx2cKO-Nes (n=3) P1 telencephalons was extracted using TRIzol (Invitrogen). Strand-specific and barcode-indexed RNA-seq libraries were generated from 1 μg total RNA each after poly-A enrichment using the Kapa Stranded mRNA-seq kit (KapaBiosystems) following manufacturer instructions. Libraries were analyzed with a Bioanalyzer 2100 instrument (Agilent), quantified by fluorometry on a Qubit instrument (Life Technologies) and pooled in equimolar ratios. The pool was quantified by qPCR with a Kapa Library Quant kit (Kapa Biosystems) and sequenced on one high output flowcell of an Illumina NextSeq 500 (Illumina) with paired-end 40 bp reads.

### RNA analyses

Total RNA from CD-1 wild type telencephalon was isolated using TRIzol. iSCRIPT system (Bio-RAD) was used for cDNA reverse transcription. Arl4c or β-actin transcripts were quantified by real-time PCR using gene-specific primers and iTaq Universal SYBR Green Supermix (Bio-Rad). ΔCt (ß-actin Ct – Arl4c-Ct) values were used to compare the Arl4c expression at the different time points. Primers used were: β-actin, 5′-CTAAGGCCAACCGTGAAAAG-3′ and 5′-ACCAGAGGCATACAGGGACA-3′; and Arl4c, 5’-AGTCTCTGCACATCGTTATGC-3’ and 5’-GGTGTTGAAGCCGATAGTGGG-3’.

### Tandem Mass Tag Mass Spectrometry

P1 Rbx2 fl/fl (n=4) and Rbx2cKO-Nes (n=3) pups were intracardially perfused with a solution of PBS and protease and phosphatase inhibitors (1mM PMSF, 10µg/ml Aprotinin, 10µg/ml Leupeptin, 10mM NaF, 1mM Na_3_VO_4_). Telencephalons from these animals were quickly dissected and snap frozen. Samples were labelled using the isobaric Tandem Mass Tags (TMT) method and analyzed and quantified by mass spectrometry. Briefly, proteins were lysed and processed for sequential digestion using LysC and trypsin. Peptides from each embryo were labelled with the TMT reagents, such the reporter ions at m/z of 126, 127N, 127C, 128N for control samples (i.e. Rbx2 fl/fl) and 128C, 129N, and 131 for Rbx2 mutant samples would be generated in the tandem spectrometry. Liquid chromatography, MS3 tandem mass spectrometry, and data analysis were carried out as previously described (McAlister et al., 2014; Ting et al., 2011).

### Western Blot Analyses

Total protein extracts from dissected cortices or hippocampi were prepared by homogenizing tissues in an appropriate volume of ice-cold lysis buffer (50mM HEPES pH7.5; 150mM NaCl; 1.5mM MgCl2; 1mM EGTA; 10% glycerol; 1% Triton X-100) using a Pellet Pestle Motor tool (Kontes). Protease and phosphatase inhibitors (1mM PMSF, 10µg/ml Aprotinin, 10µg/ml Leupeptin, 10mM NaF, 1mM Na_3_VO_4_) were added to the lysis buffer just prior to homogenization. The homogenate was cold-centrifuged for 15 min at 14,000 rpm. 20µl of cleared supernatant were denatured at 95C for 10 min in sample buffer (10mM Tris pH 6.5; 150mM β-mercaptoethanol; 0.5%SDS, 5% glycerol and 0.0125% bromophenol blue) and loaded into a (SDS)-polyacrylamide gel, to finally be transferred to a 0.2µm nitrocellulose membrane (Bio-RAD) using standard methods. Membranes were blocked for 1 hour with blocking solution (Tris buffered saline with 0.1% Tween-20 (TBS-T), 5% non-fat milk powder or 5% BSA), washed with TBS-T and incubated overnight with primary antibodies (anti-Actin 1:10000 Santa Cruz Biotechnology #sc1616; anti-Dab1 1:2000 Sigma-Aldrich #HPA052033, anti-Fyn 1:1000 Santa Cruz Biotechnology #sc-16; anti-Tuj1 1:20000 BioLegend #801201; anti-HA 1:10000 BioLegend #901501; anti-Arl4c 1:1000 Proteintech #10202-1; anti-Cytohesin-1 1:1000 Fisher Scientifics #MA1060). Specific HRP-conjugated secondary antibodies were incubated for 45 min at room temperature in blocking solution. Proteins were identified using SuperSignal West Pico ECL substrate (ThermoFisher Scientific) or ProSignal Femto ECL reagent (Genesee Scientific) by exposing to X-ray films. Films were scanned on an Epson perfection V600 photo scanner and converted to 16-bit images for band densitometry quantification using FIJI software (ImageJ2 v1.51t, NIH). Protein abundancy was calculated upon band densitometry analysis compared to littermate controls and results were normalized using the detection of Actin or Tuj1 as a loading control. All images were assembled using Adobe Photoshop and Illustrator.

### Hippocampal quantifications and Statistical Methods

For all *in vivo* data sets biological replicates were obtained from at least two different litters or *in utero* electroporations. Specific number of biological replicates (Ns) are specified in each figure/figure legend. For hippocampal size and DKK3+ cell quantification, we quantified the hippocampus in each hemispheres of three sections containing the dorsal hippocampus in which the habenula was visible. For IUEs, all detected GFP+ cell bodies in the CA1/CA2 regions were quantified for at least three electroporated sections in each brain; for dendrite quantifications only those cells where a clear apical dendrite(s) were observed were used for our quantifications. For *in vitro* Arl4c puncta analysis, five neurons were quantified per embryo and compared to wild type littermate controls. Unless indicated, histograms represent the mean ± the standard error of the mean (SEM). Specific statistical tests used in each case are indicated in figure legends. All statistical analyses and plot generation were performed using Prism 7 (GraphPad).

## Acknowledgements

We thank Dr. Fang-Jen Lee and Dr. Thomas Roberts for their generosity sharing reagents; Dr. Jonathan A. Cooper for critical reading of the manuscript; Dr. Monica Britton for her help analyzing RNA sequencing data; and Ysidra Camarena for technical assistance. We particularly thank Dr. Ryan C. Kunz and the Thermo Fisher Scientific Center for Multiplexed Proteomics service core (Harvard University) for running our TMT-MS experiment. This work was supported by NIH grants R01 NS109176 to S.S.; and R21 NS101450 to S.S. and A.L.T.. We also benefited from the use of the National Eye Institute Core Facilities [supported by P30 EY012576].

## Author Contribution

J.S.H., A.L.T and S.S. conceived and designed the experiments. J.S.H. performed all neuronal cultures, Sholl analyses, western blotting, and quantitative RT-PCR experiments. K.H. and S.S. did in utero electroporations and tissue processing. A.M.M. and A.L.T performed and analyzed the RNA sequencing experiments. J.S.H., K.H., R.V.R., C.P.C., Y.H., J.S., C.Y.C performed histological studies, imaging, and data analysis. S.S. wrote the manuscript with input from all the authors.

