## Supplemental Figures for "CRL5-dependent regulation of Arl4c and Arf6 controls hippocampal morphogenesis"

Supplementary Figure 1

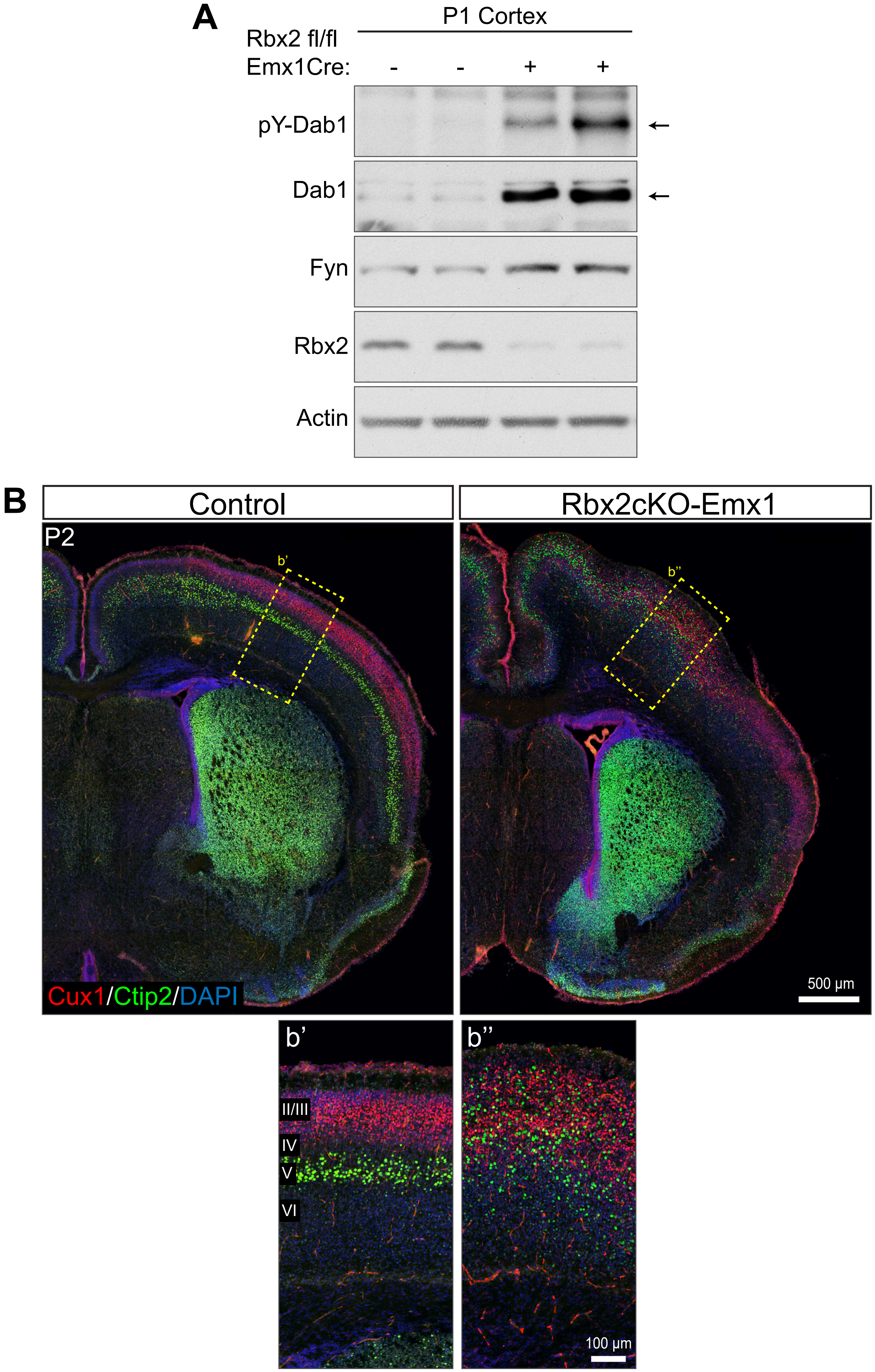

Supplementary Figure 2

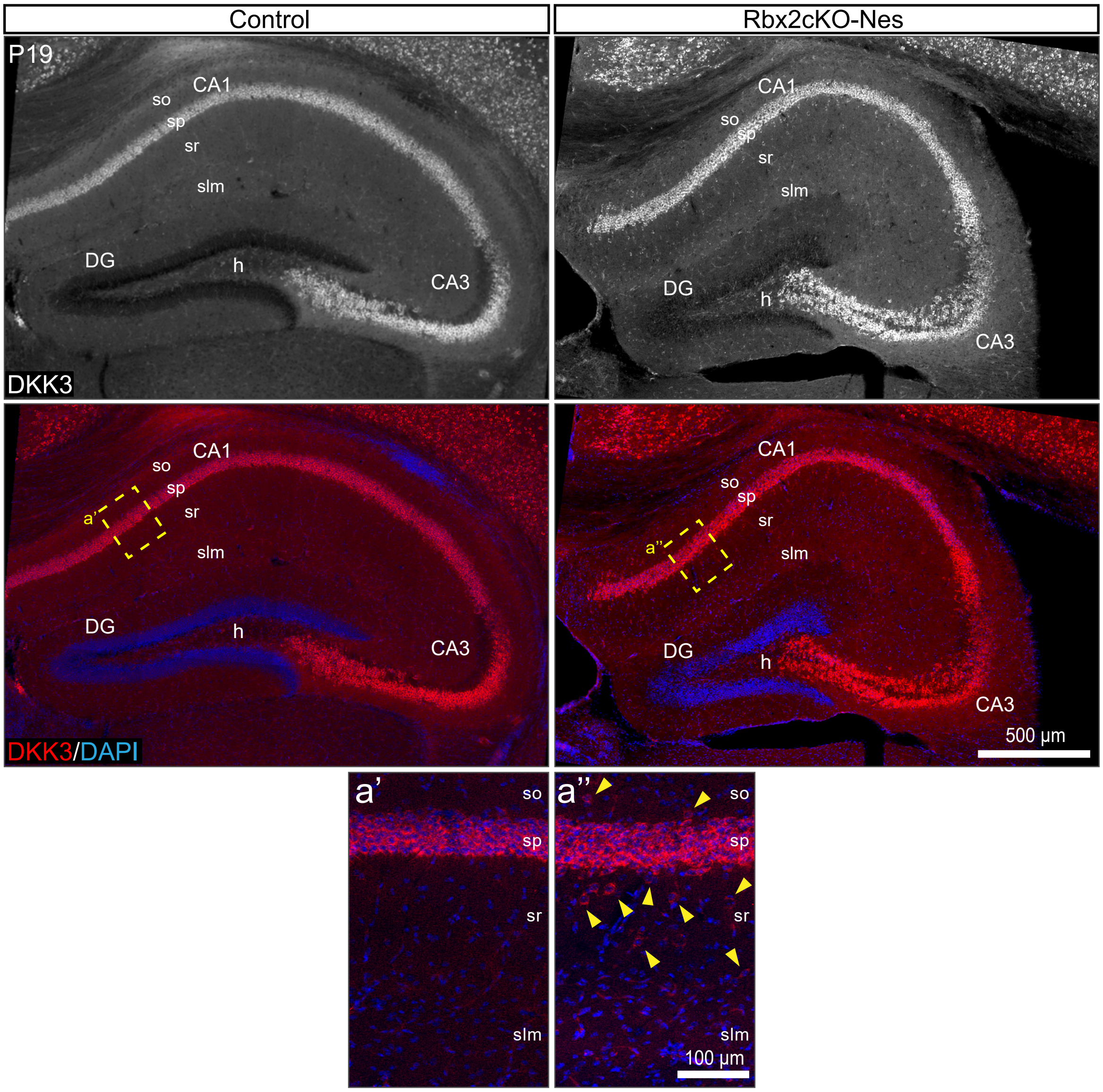

Supplementary Figure 3

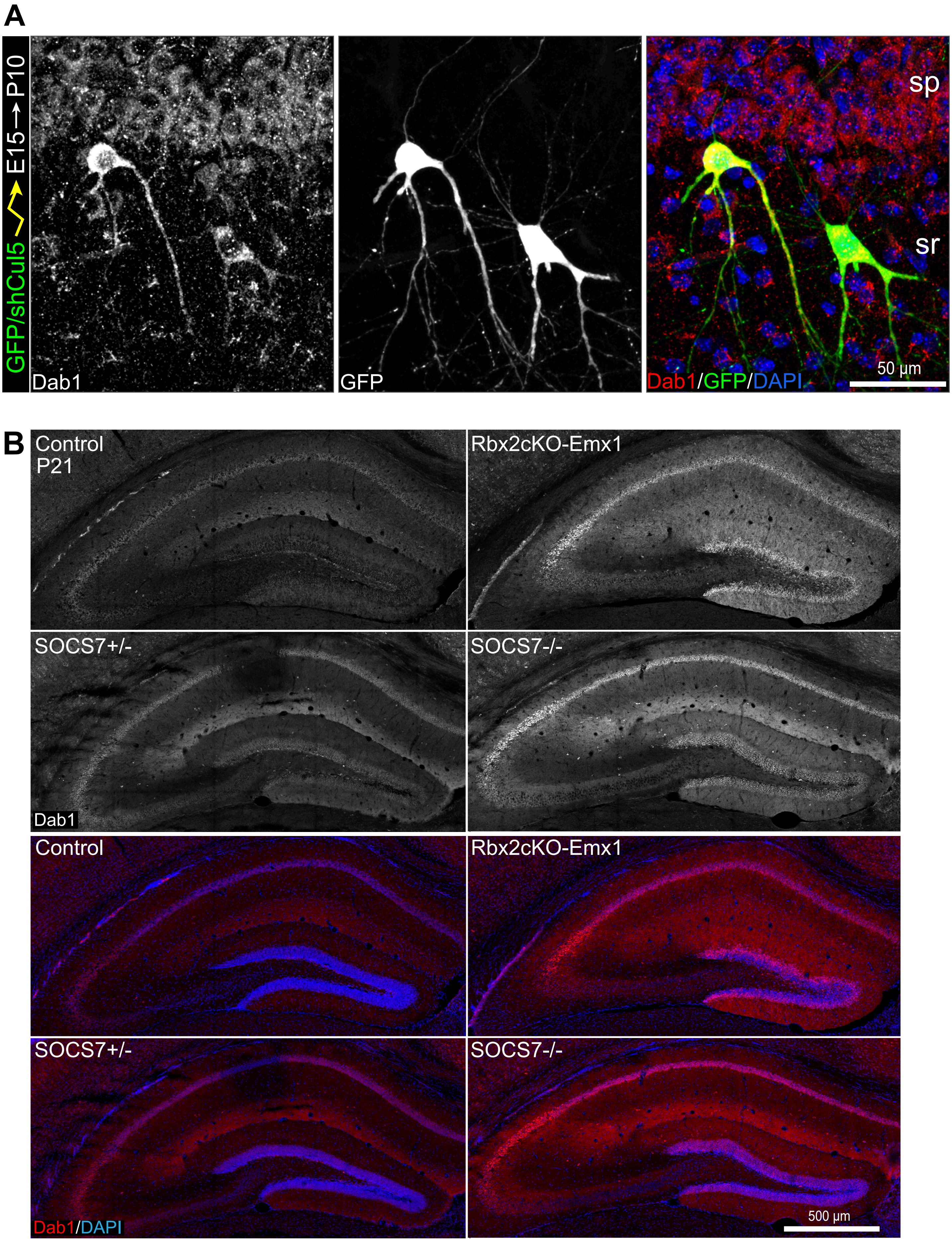

Supplementary Figure 4

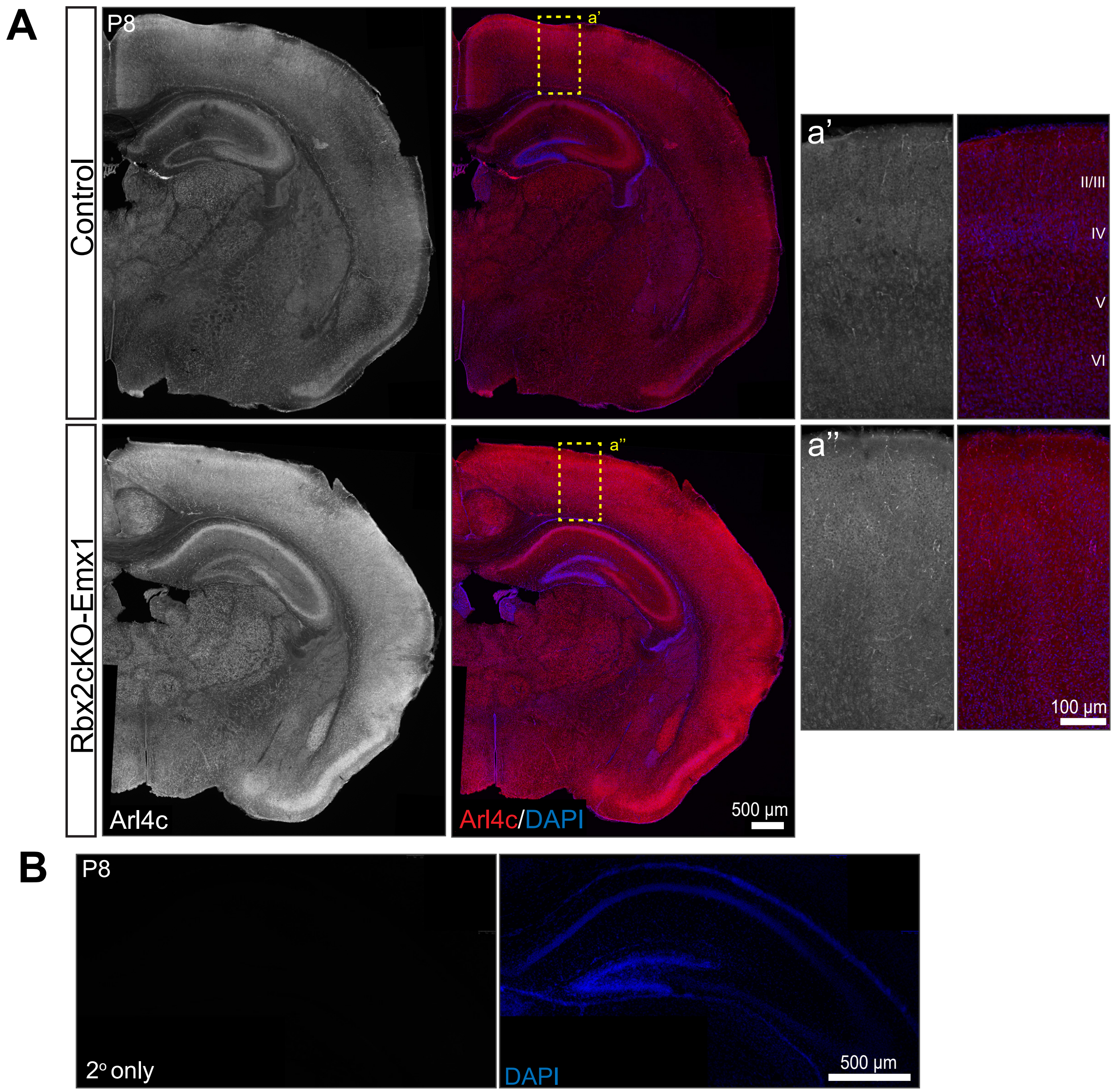

Supplementary Figure 5

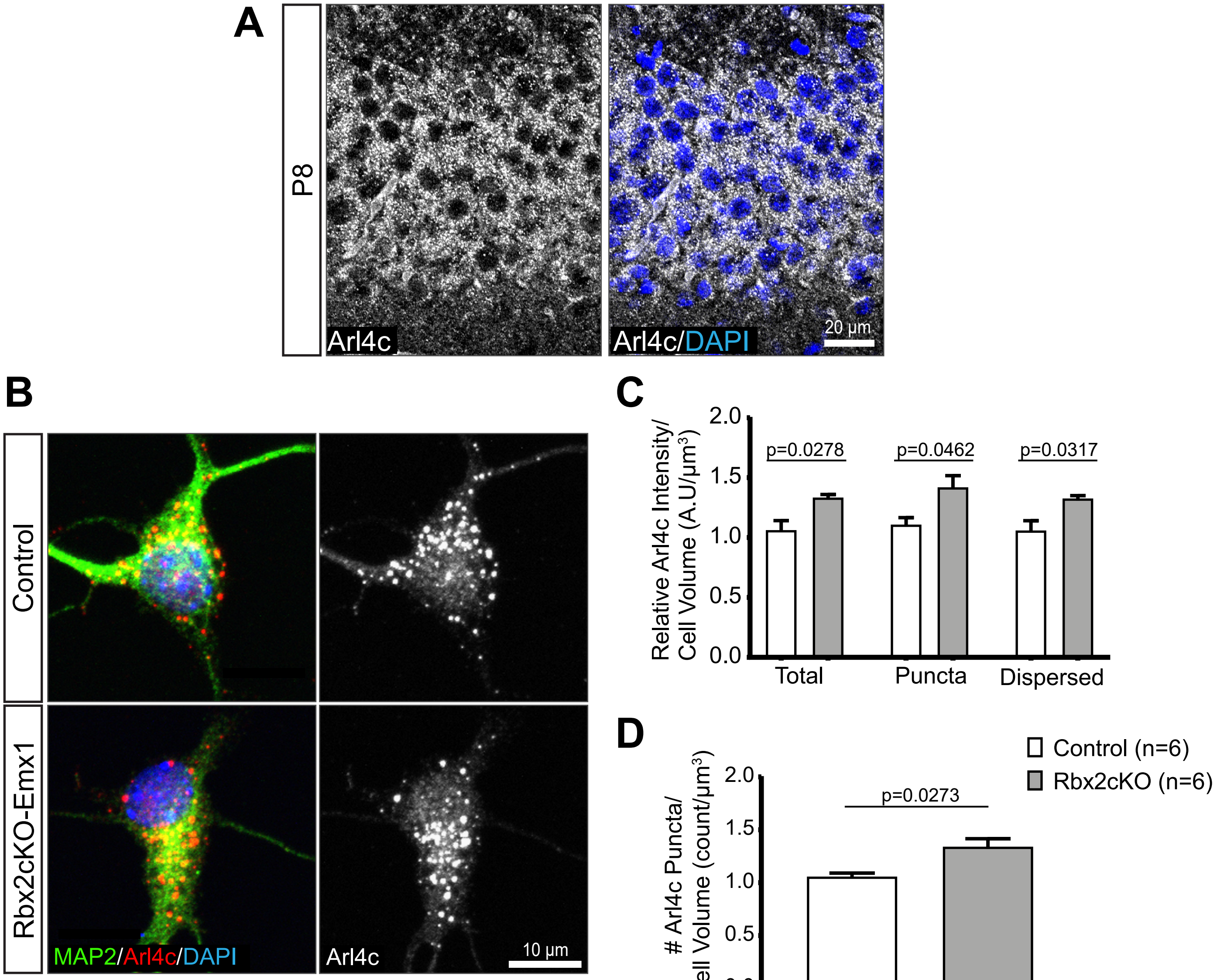

Supplementary Figure 6

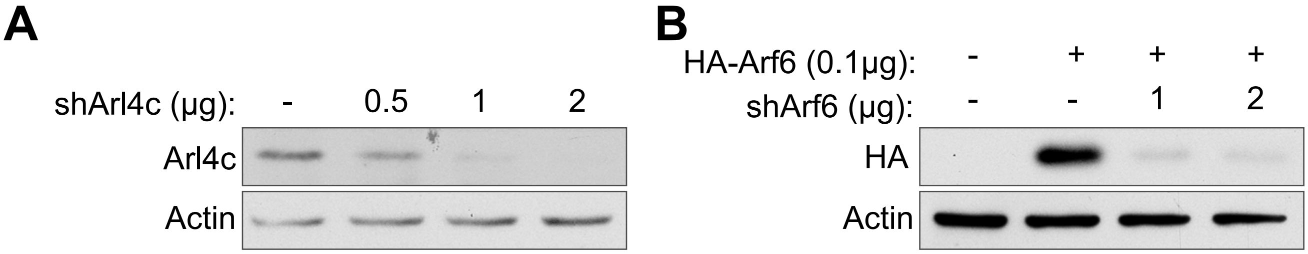
