## Supplemental Figure legend for "CRL5-dependent regulation of Arl4c and Arf6 controls hippocampal morphogenesis"

**Figure S1. Rbx2cKO-Emx1 cortex accumulates pY-Dab1, Dab1 and Fyn and shows layering defects similar to Rbx2cKO-Nes cortex.**

(A) Cortical lysates of Rbx2 fl/fl and Rbx2cKO-Emx1 at P1 were analyzed by western blotting using antibodies against p-Tyr, Dab1, Fyn, Rbx2 and Actin. Rbx2cKO-Emx1 showed accumulation of pY-Dab1, Dab1 and Fyn compared to their littermate controls.

(B) Coronal sections of P2 Rbx2 fl/fl (Control) and Rbx2cKO-Emx1 brains were stained with anti-Cux1 and anti-Ctip2 antibodies and counterstained with DAPI. PNs show dispersion in the neocortex as seen in the Rbx2cKO-Nes neocortex (Simo and Cooper, 2013). (b'-b'') High-magnification images of yellow box in (B).

**Figure S2. Rbx2cKO-Nes hippocampal PNs are ectopically localized in *so*, *sr*, and *slm*.**

Coronal sections of P19 Rbx2 fl/fl (Control) and Rbx2cKO-Nes brains were stained with anti-DKK3 antibodies and counterstained with DAPI. PNs were observed in *so*, *sr*, *slm* in Rbx2cKO-Nes as observed in the Rbx2cKO-Emx1 hippocampus (related to Figure 1). (a'-a'') Magnified images of CA1 region. Yellow arrowheads indicate mispositioned DKK3+ PNs.

**Figure S3. Dab1 accumulation is not sufficient to cause layering defect observed with CRL5 inactivity.**

(A) GFP/shCul5 electroporated brains were sectioned and stained with an antibody against Dab1 and counterstained with DAPI. Both GFP/shCul5+ PNs show over-migration, abnormal apical dendrites, and Dab1 accumulation in comparison to unelectroporated PNs in the *sp*.

(B) Coronal sections of P21 control (Rbx2 fl/fl), Rbx2cKO-Emx1, SOCS7+/-, and SOCS7-/- brains were stained with an anti-Dab1 antibody and counterstained with DAPI. Both Rbx2cKO-Emx1 and SOCS7-/- hippocampi accumulate Dab1 at similar levels and in similar areas.

**Figure S4. Arl4c has high expression in the telencephalon and accumulates in Rbx2cKO-Emx1 brains.**

(A) Coronal sections of P8 Rbx2 fl/fl (Control) and Rbx2cKO-Emx1 brains were stained with an anti-Arl4c antibody and counterstained with DAPI. In control brains, Arl4c is widely expressed in the brain, with a stronger expression in the neocortex and the hippocampus. (a'-a'') High-magnification images of the neocortex in (A, yellow box).

(B) Negative control for Arl4c antibody staining. Brain sections were process in parallel with samples in (A) but Arl4c antibody was excluded.

**Figure S5. Arl4c localizes at the plasma membrane and in cytoplasmic puncta in pyramidal neurons and Rbx2-depletion increases cytoplasmic puncta *in vitro.***

(A) Coronal sections of P8 wild type hippocampus stained with an anti-Arl4c antibody and counter stained with DAPI. Representative image of hippocampal *sp* shows Arl4c punctum *in vivo.*

(B) Hippocampal primary cultures were fixed after 4 days in culture and stained with antibodies against MAP2 and Arl4c and counterstained with DAPI. Arl4c localizes to puncta throughout the cell body and in neurites in both Rbx2 fl/fl (Control) and Rbx2cKO-Emx1 hippocampal neurons.

(C-D) Arl4c signal intensity (C) and number of Arl4c puncta (D) in Control and Rbx2cKO-Emx1 neurons were quantified. Five neurons per biological replicate (n=6) from at least three different litters were quantified and normalized to cell volume and to control littermates. Rbx2cKO-Emx1 neurons accumulate Arl4c (C) and have increased number of Arl4c+ puncta (D). Statistics, Student's t-test.

**Figure S6. shArl4c and shArf6 show high knockdown efficiency in HEK293T cells.**

(A) HEK293T cells were transfected with shArl4c expressing plasmids in varying amounts. Cells were lysed after 24 hours and endogenous Arl4c levels were observed by western blotting using an anti-Arl4c antibody. Decreased levels of Arl4c were observed in shArl4c concentration-dependent fashion.

(B) HEK293T cells were co-transfected with HA-Arf6 and shArf6 expressing plasmids at indicated amounts and lysed after 24 hours for western blotting analysis. Arf6 levels were observed using antibody against the HA tag. shArf6 expressing samples show significant decrease in HA signal compared to control.

Simo, S., and Cooper, J.A. (2013). Rbx2 regulates neuronal migration through different cullin 5-RING ligase adaptors. Dev Cell *27*, 399-411.
